# The WPRE improves genetic engineering with site-specific nucleases

**DOI:** 10.1101/126904

**Authors:** Jessica M. Ong, Christopher R. Brown, Matthew C. Mendel, Gregory J. Cost

## Abstract

**Abstract:** Inclusion of the woodchuck hepatitis virus post-transcriptional response element (WPRE) in the 3’ UTR of mRNA encoding zinc-finger or TALE nucleases results in up to a fifty-fold increase in nuclease expression and a several-fold increase in nuclease-modified chromosomes. Significantly, this increase is additive with the enhancement generated by transient hypothermic shock. The WPRE-mediated improvement is seen across several types of human and mouse primary and transformed cells and is translatable *in vivo* to the mouse liver.

---

The woodchuck hepatitis virus post-transcriptional response element (WPRE) is thought to promote nuclear export of viral genomic mRNA (1). As a result, the WPRE is often included in viral gene transfer vectors where it can improve viral titer and subsequent transgene expression (2). The effect of the WPRE is idiosyncratic, improving expression of some transgenes in some cell types but occasionally having neutral or even negative effects (3, 4, 5, 6, 7, 8, 9).

There are conflicting reports as to how the WPRE works. In some contexts its presence increases steady-state mRNA levels (10) while in others the WPRE seems to improve transcriptional termination, having no effect on nuclear export or mRNA half-life (11). As there is one claim of an improvement in cytoplasmic mRNA metabolism resulting in increased translation (12), we investigated whether inclusion of the WPRE in site-specific nuclease mRNA would improve nuclease expression and genome modification.

To determine whether inclusion of the WPRE in the 3’ UTR of a nuclease transcript would improve non-homologous end joining (NHEJ) DNA repair, we transfected *in vitro* transcribed mRNAs encoding *CCR5*-specific zinc-finger nucleases (ZFNs) either containing the WPRE or not and assayed the *CCR5* locus by high-throughput DNA sequencing. The presence of a WPRE increased the frequency of NHEJ by two‐ to ten-fold in CD34-positive hematopoietic stem and progenitor cells (HSPCs), CD4-positive T cells, and CD8-positive T cells (Figure 1a, b, c). An increase in mutagenic NHEJ was seen when the WPRE was included in *CCR5* TALE nuclease mRNAs as well (Figure 1d). We quantitated ZFN protein levels in transfected K562 cells and found that the WPRE-containing mRNAs yielded a steady-state level of ZFN protein between four‐ and 67-fold higher depending on the dose transfected (Figure 2) with improvements in NHEJ similar to primary cells (data not shown).

**Figure 1.**
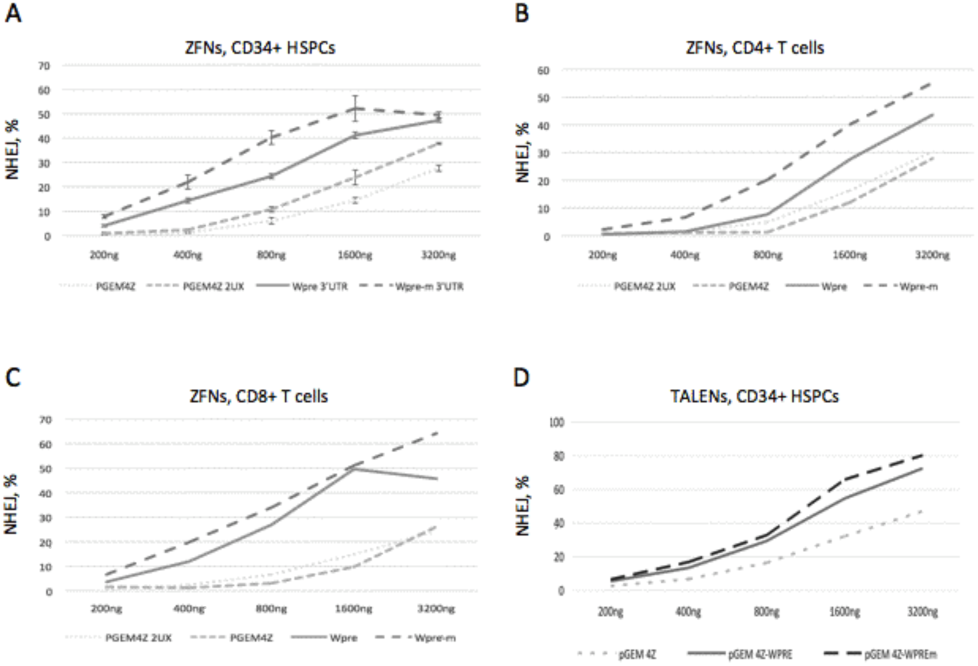
NHEJ is enhanced when the WPRE is included in the 3’ UTR. NHEJ from transfection in triplicate of 200, 400, 800, 1600, or 3200 ng of mRNA encoding the human CCR5-specific ZFNs into CD34-positive HSPCs (A), CD4+ T cells (B), and CD8+ T cells (C). NHEJ from transfection in triplicate of 200, 400, 800, 1600, or 3200 ng of mRNA encoding human CCR5-specific TALENs into CD34-positive HSPCs (D). The nuclease mRNAs contained either no addition to the 3’ UTR (pGEM4X), a dimer of an RNA stability element from the *X. laevis* beta-globin gene (pGEM4Z 2UX), a wild-type J04514 WPRE (WPRE), or a mutant J02442 WPRE (WPRE-m).

**Figure 2.**
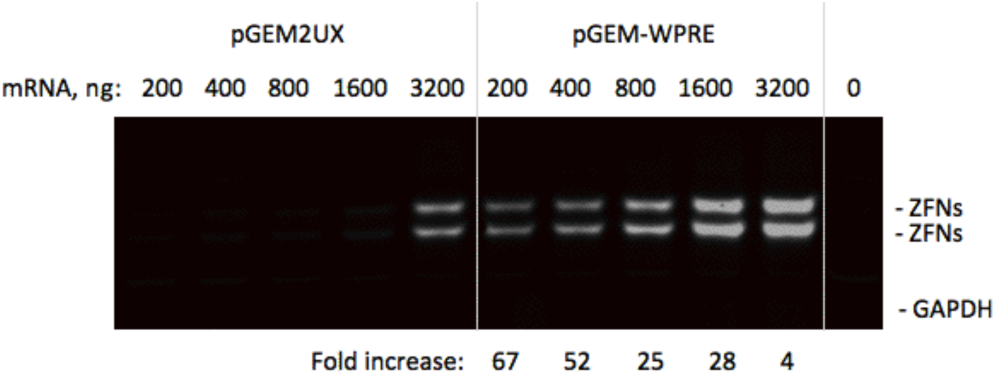
Increased ZFN protein after transfection of 200, 400, 800, 1600, or 3200 ng of mRNA encoding the human CCR5 ZFNs into K562 cells. The ZFN mRNAs contained either the above 2UX element or a wild-type J04514 WPRE (WPRE). The fold increase in ZFN protein is shown below the gel.

A brief hypothermic shock has also been shown to increase NHEJ activity. To test whether hypothermia and the WPRE effect increased NHEJ via the same mechanism, we individually transfected CD34-positive HSPCs with mRNA encoding three separate ZFN pairs either with or without the WPRE in their 3’ UTRs. The transfected cells were divided in half, with one half incubated at 30° C for 18 hours and the other half maintained at 37° C. At *AAVS1, IL2Rγ*, and *CCR5*, the WPRE-mediated improvement in NHEJ was additive with the improvement gained from hypothermic shock (Figure 3). We conclude that hypothermic shock and the WPRE function via independent and complementary biological pathways.

**Figure 3.**
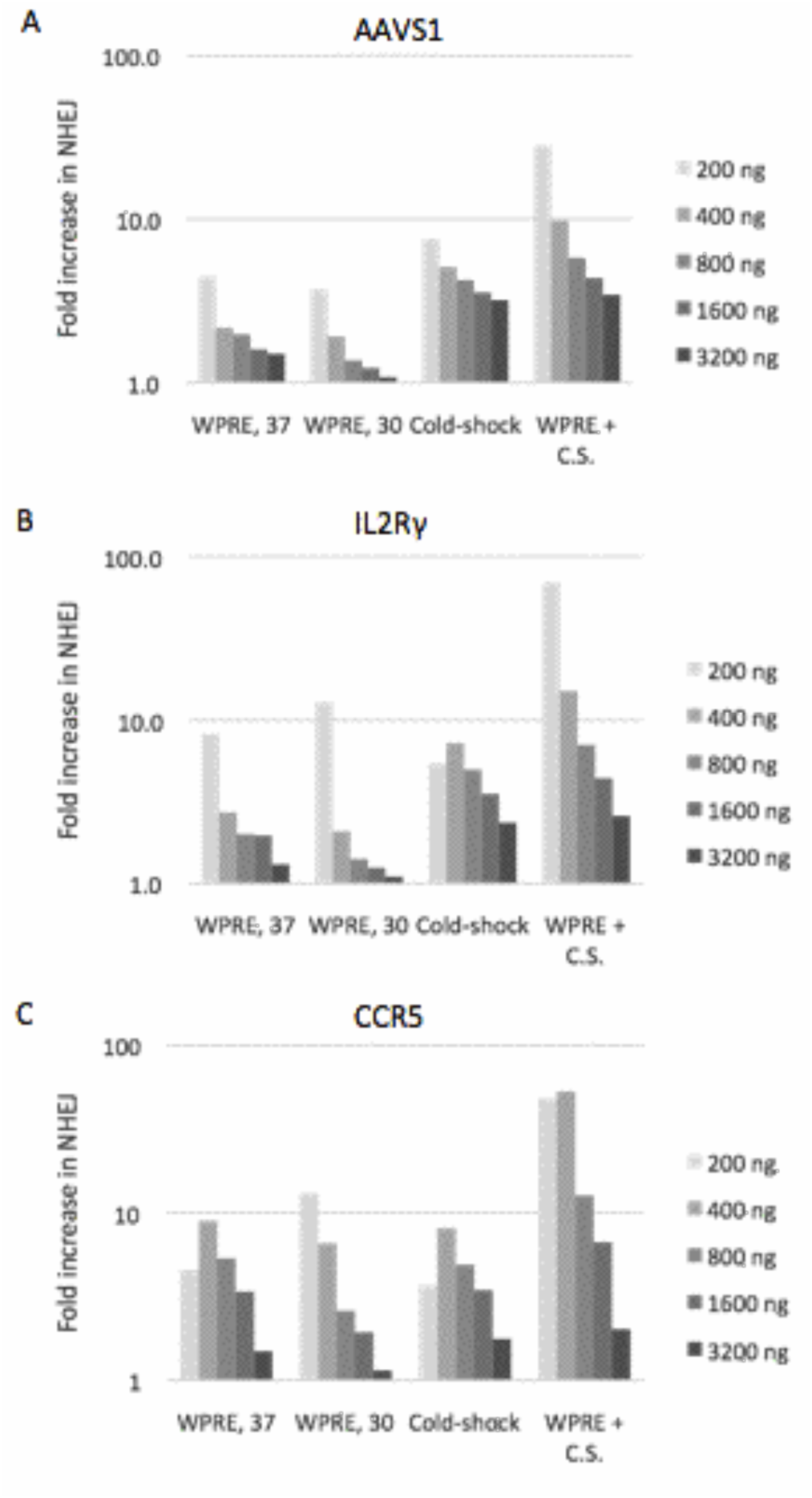
The fold improvement in NHEJ achieved by different doses of three different ZFN pairs under four experimental conditions: i) WPRE and culture at 37° C (WPRE, 37; compared to no WPRE and 37° C culture); ii) WPRE and culture at 30° C (WPRE, 30; compared to no WPRE and 30° C culture); iii) hypothermic shock with no WPRE (Cold-shock; compared to no WPRE and 37° C culture); iv) the combined effect of the WPRE and hypothermic shock (WPRE + C.S.; compared to no WPRE and culture at 37°C). Logarithmic scale.

**Figure 4.**
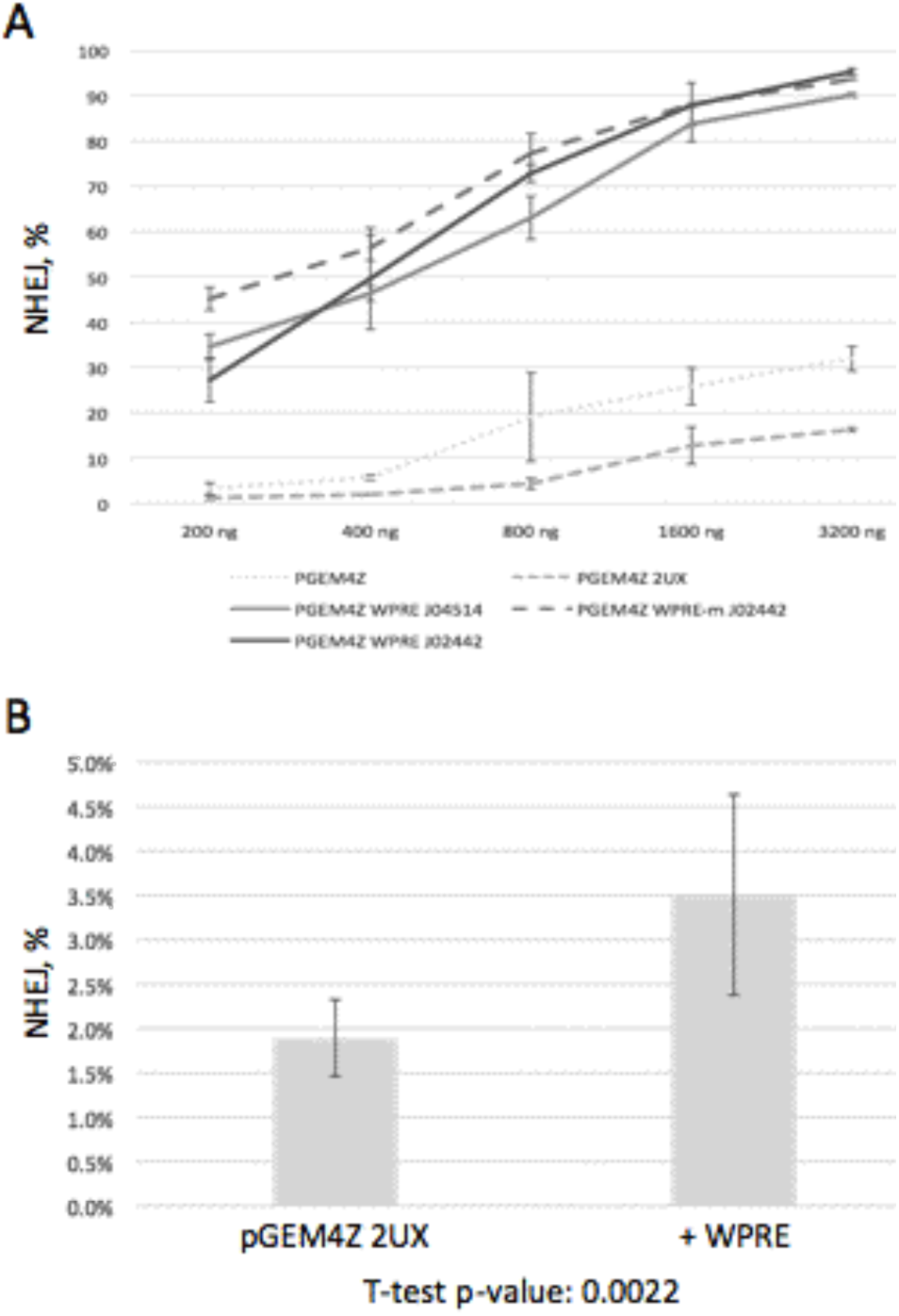
WPRE-mediated enhancement of NHEJ in mouse cells and in mice. A) NHEJ DNA repair after transfection in triplicate of 25, 50, 100, 200, or 400 ng of mRNA encoding the mouse Albumin-specific ZFNs into Hepa1-6 cells. The ZFN mRNAs contained either no addition to the 3’ UTR (pGEM4X), the 2UX element (pGEM4Z 2UX), a wild-type WPRE (pGEM4Z WPRE J04514 and pGEM4Z WPRE J02442), or a mutant J02442 WPRE (pGEM4Z WPRE-m J02442). B) NHEJ DNA repair in the mouse liver (N=6; measured in triplicate in each mouse) after lipid nanoparticle delivery of the same mRNA as in part A. A Student’s two-tailed T-test with two-sample equal variance indicated that the improvement seen with the addition of the WPRE was highly unlikely to be due to chance (p=0.0022)

Use of the WPRE produces meaningful increases in NHEJ in cells treated *ex vivo*, including mouse cells (Figure 2a). As mRNA delivery to primary cells *in vivo* is a promising mode of therapeutic treatment we examined the effect of the WPRE when lipid nanoparticle-formulated ZFN mRNA was delivered intravenously to mice. In contrast to viral delivery of WPRE-containing transgenes to the mouse liver (4), delivery of WPRE-containing mRNA resulted in NHEJ at a frequency nearly double that of a non-WPRE-containing transcript (Fig. 2b).

We find that inclusion of the WPRE in the 3’ UTR of ZFN and TALE nuclease mRNA constructs dramatically boosts nuclease expression and the percentage of chromosomes cleaved by the nuclease. In addition to the ZFN pairs shown here, similar results have been obtained with four additional pairs (data not shown).

## Discussion

Can the level of gene editing achieved with WPRE-containing mRNAs be obtained simply by transfection of additional conventional mRNA? In human cells the WPRE increases the specific activity of the mRNA several fold but in most cases does not increase the maximal level of NHEJ. In contrast, data from mouse cells suggests that the plateau level of NHEJ is higher with use of the WPRE (Fig 2b). Simply using less mRNA confers significant advantages regardless of cell type as it reduces the consequences from the induction of the innate immune system.

Concerns have been raised about the possibility – however slight – that the WPRE might promote cellular transformation and neoplasia due to expression of a truncated woodchuck hepatitis virus X protein from within the WPRE. Delivery of a WPRE-containing mRNA completely eliminates the promoter and prevents X protein mRNA production. Further, a WPRE containing mutations that prevent X protein expression (13) was equally or more effective at stimulating NHEJ and could be used to prevent X protein production from rare translational re-initiation events. Finally, transfection of WPRE-containing mRNA indicates that the WPRE need not act during transcription or transcriptional termination to improve transgene expression.

## Methods

### Cell culture, transfection and DNA analysis

HSPCs were grown in X-VIVO 10 with STEMSPAN CC110 cytokines plus 20 ug/mL IL-6 and transfected in triplicate with a 96-well BTX ECM 830 transfection device using BTXpress electroporation solution with mode LV, 250V, and a 4 ms pulse length. CD8-positive and CD4-positive T cells were grown in X-VIVO 15 with 10% FBS and 100 ug/mL IL-2, stimulated with anti-CD3/CD28 Dynabeads at 10∧8/mL, and transfected with a 96-well BTX ECM 830 transfection device using BTXpress electroporation solution with mode LV, 250V, and a 4 ms pulse length. K562 cells were grown in Iscove’s Modified Dulbecco’s Medium with 10% fetal bovine serum and transfected with a Lonza Amaxa nucleofector shuttle using solution SF and program FF-120. Hepa1-6 cells were grown in Dulbecco’s Modified Eagle’s Medium with 10% fetal bovine serum and transfected in triplicate with a Lonza Amaxa nucleofector shuttle using solution SG and program DS-150. Genomic DNA was harvested, amplified by PCR and ∼15,000 alleles per sample analyzed on an Illumina MiSeq high-throughput DNA sequencer.

### Western blotting

Protein lysates were analyzed by Western blot using a mouse anti-FLAG IgG1 primary antibody and with a mouse anti-GAPDH IgM primary antibody. Secondary goat anti-mouse IgG1-800 nm fluor and IgM-680 nm fluor antibodies were then applied followed by interrogation of the membrane on a LiCor Odyssey CLx. The ZFN signal was normalized by the GAPDH signal to account for differences in the total amount of protein loaded on the gel.

### DNA sequences, plasmid construction, and in vitro transcription

The wild-type WPRE sequences used are found in Genbank files J04514 and J02442; the mutant WPRE sequence is from Zanta-Bousiff *et al*. The WPRE was PCR amplified using the primers 5’-gta agt cta gaa atc aac ctc tgg att aca aaa t-3’ and 5’-atg aac atc tag aca ggc ggg gag gcg gc-3’, the amplicon digested with XbaI and cloned in the forward orientation into the XbaI site of a pGEM-based plasmid.

The *CCR5* ZFNs are SBS 8266 and SBS 8196z; the *CCR5* TALE nucleases are SBS 101041 and SBS 101047 with C63 truncation points; the mouse Albumin ZFNs are SBS 48641 and SBS 31523.

Plasmids were linearized for *in vitro* transcription with SpeI (CCR5, AAVS1, and Albumin ZFNs), EagI (CCR5 TALE nucleases). mRNA for transfection was synthesized with an Ambion mMessage mMachine kit according to the manufacturer’s recommendations. mRNA for *in vivo* delivery was manufactured by TriLink Inc. LNPs were manufactured by Acuitas Therapeutics Inc.

## References

1. Donello JE et al. Woodchuck hepatitis virus contains a tripartite posttranscriptional regulatory element. J Virol. 1998 Jun;72(6):5085–92.

2. Zufferey R et al. Woodchuck hepatitis virus posttranscriptional regulatory element enhances expression of transgenes delivered by retroviral vectors. J Virol. 1999 Apr;73(4):2886–92.

3. Breckpot K et al. Lentivirally transduced dendritic cells as a tool for cancer immunotherapy. J Gene Med. 2003 Aug;5(8):654–67.

4. Cotugno G et al. Impact of age at administration, lysosomal storage, and transgene regulatory elements on AAV2/8-mediated rat liver transduction. PLoS One. 2012;7(3):e33286.

5. Klein R et al. WPRE-mediated enhancement of gene expression is promoter and cell line specific. Gene. 2006 May 10;372:153–61.

6. Kraunus J et al. Self-inactivating retroviral vectors with improved RNA processing. Gene Therapy 2004 Nov;11(21):1568–78.

7. Mangeot PE et al. High levels of transduction of human dendritic cells with optimized SIV vectors. Molecular Therapy 2002 Mar;5(3):283–90.

8. Schambach A et al. Context dependence of different modules for posttranscriptional enhancement of gene expression from retroviral vectors. Molecular Therapy 2000 Nov;2(5):435–45.

9. Werner M, Kraunus J, Baum C, Brocker T. B-cell-specific transgene expression using a self-inactivating retroviral vector with human CD19 promoter and viral post-transcriptional regulatory element. Gene Therapy 2004 Jun;11(12):992–1000.

10. Loeb JE et al. Enhanced expression of transgenes from adeno-associated virus vectors with the woodchuck hepatitis virus posttranscriptional regulatory element: implications for gene therapy. Hum Gene Therapy 1999 Sep 20;10(14):2295–305.

11. Higashimoto T et al. The woodchuck hepatitis virus post-transcriptional regulatory element reduces readthrough transcription from retroviral vectors. Gene Therapy 2007 Sep;14(17):1298–304.

12. Loeb JE. Ph.D. Thesis, Univeristy of California, San Diego. 2000.

13. Zanta-Boussif MA et al. Validation of a mutated PRE sequence allowing high and sustained transgene expression while abrogating WHV-X protein synthesis: application to the gene therapy of WAS. Gene Therapy 2009 May;16(5):605–19.

